# Zyxin Restricts Viral Fusion and Entry across Multiple Virus Families

**DOI:** 10.1101/2025.10.03.680228

**Authors:** Qing Fan, Jenai Quan, Gregory A. Smith, Richard Longnecker

## Abstract

The entry of enveloped viruses into host cells requires fusion of the viral envelope with a host cell membrane. The composition and dynamics of cellular membranes are impacted by the underlying cytoskeleton. Zyxin, a cytosolic protein that bridges the actin cytoskeleton with adhesion receptor proteins, was recently identified as an antiviral factor that antagonizes HSV-1 entry. To determine if zyxin specifically interferes with membrane fusion, we examined its impact in a quantitative cell-cell fusion assay. HSV-1 entry proteins exhibited enhanced fusion activity with retinal pigment epithelial (RPE) cells when the cells were knocked out (KO) for zyxin. Zyxin-KO cells also showed enhanced fusion activity with pseudorabies virus (PRV), paramyxovirus and rhabdovirus fusion proteins. Additionally, the size of plaques formed following infection with members of each viral family were increased in the absence of zyxin. Bulk RNA sequencing of wild-type (WT) and zyxin-KO RPE cells identified 18 genes that enrich into several ontology groups of interest, including regulation of membrane potential, herpes simplex virus 1 infection (CCL2 and ZNF14), NABA core matrisome (CCL2 and ZNF14), extracellular matrix organization (CTSK, SERPINE1 and TGFB2), cell-cell adhesion (RAC2 and TGFB2), anchoring fibril formation, and regulation of the MAPK cascade (ACKR3 and TGFB2). These results highlight zyxin as a broadly active antiviral factor and suggest potential therapeutic implications for viral infections.

**IMPORTANCE:** Enveloped viruses enter host cells by fusing their envelope with a cellular membrane. This process is triggered when a viral glycoprotein(s) engages with a cellular membrane receptor. Less well understood are the cytoplasmic factors that indirectly govern viral fusion and entry. We report that zyxin, a cellular protein that bridges cell-adhesion receptors with the underlying actin cytoskeleton, antagonizes the fusion and entry of several enveloped viruses.

## INTRODUCTION

Herpes simplex virus 1 (HSV-1) infects humans and causes recurrent mucocutaneous lesions on the mouth, face, or genitalia. On rare occasions, infections progress to meningitis or encephalitis (1). The entry of HSV-1 into host cells requires the coordinated action of four viral glycoproteins: gB, gD, gH, and gL (2–5). The binding of gD to one of several host receptors results in a conformational change that triggers the fusogenic activity of gB via gH/gL (4–6). Entry receptors that bind to gD include herpes virus entry mediator (HVEM) (7), nectin-1 (8, 9), nectin-2 (10, 11), and modified heparan sulfate (12, 13). Another receptor, paired immunoglobulin-like type 2 receptor (PILRα), binds to gB directly and enhances viral entry (14). Zyxin is a component of focal adhesions that modulates cellular mechanics by interfacing cell adhesion receptors with the underlying actin cytoskeleton and also participates in signal transduction to the nucleus (15–20). Zyxin is implicated in cancer progression, morphogenetic cell movements, gene expression, and pattern formation during embryogenesis (21–23), and recently was implicated as a protective factor to viral infection (24). In the latter study, pseudorabies virus (PRV), a veterinary herpesvirus, expressing the BioID2 proximity-labeling reporter fused to either the pUL36 or pUL37 tegument protein identified zyxin as a candidate protein encountered by incoming virus. CRISPR-Cas9 knockout of zyxin from hTERT-immortalized retinal pigmented epithelial (RPE) cells resulted in increased susceptibility to PRV and HSV-1 (24).

The initial goal of the current study was to determine whether zyxin affects cell fusion activity of HSV-1. Using an established cell-cell fusion assay that we adopted for use with RPE cells, we found that zyxin knockout (KO) RPE cells were hyperfusogenic for HSV-1 glycoproteins relative to the wild-type (WT) RPE controls. The increase in fusion was also noted with the equivalent PRV glycoproteins and with glycoproteins of a paramyxovirus and rhabdovirus. During infection, viruses grown on the KO RPE cells formed larger plaques than those produced on wild-type RPE cells. Finally, RNA sequencing of WT and KO RPE cells identified several transcripts displaying zyxin dependence and possibly implicating endocytosis as factor in dictating broad-spectrum resistance to fusion based entry of enveloped viruses.

## RESULTS

### Cell-cell fusion activities between CHO-K1 and RPE cells

Zyxin antagonizes an early step of HSV-1 and PRV infection (24). To examine whether zyxin specifically constrains virus-host membrane fusion, we first determined whether RPE cells could be used in our established cell-cell fusion assay (25). This reductionist model uses effector cells that express viral entry proteins and the T7 polymerase, and target cells that express cell entry receptors along with a T7-driven luciferase reporter. Luciferase expression is used to quantify cell-cell fusion.

Consistent with previous reports, using CHO-K1 cells as both effector and target cells (the former expressing HSV-1 gB, gD, and gH/gL), fusion activity was detected with target cells expressing either nectin-1 or PILRα (the latter showing approximately 10% fusion activity relative to nectin-1; *P* < 0.01) (Fig. 1A) (26, 27). Unfortunately, no fusion activity was detected when CHO-K1 effector and target cells were both substituted with corresponding RPE cells. The raw luciferase activity (Relative Light Units, RLU) was lower than that of the empty vector control (below 2000 RLU), corresponding to only ∼30% of the empty vector RLU (Fig. 1B). This finding indicates that fusion does not readily occur between RPE cells when the effector cells expressed HSV-1 entry glycoproteins and the target cells expressed nectin-1. Because transfected CHO-K1 were previously established to express HSV-1 glycoproteins at the cell surface, we next used CHO-K1 cells as effector cells in combination with RPE target cells. In this setup, fusion activity was detected with WT RPE, zyxin-KO RPE, and rescued RPE cells (zyxin-eGFP) expressing either nectin-1 or PILRα. Notably, the fusion activity in zyxin-KO cells was higher than that in the WT and rescue cells (*P* < 0.01). Furthermore, to our surprise, fusion with PILRα-expressing RPE targets was as efficient as the Nectin-1 RPE targets (Fig. 1C), a result we had not previously observed when using CHO cells only.

**Fig. 1.**
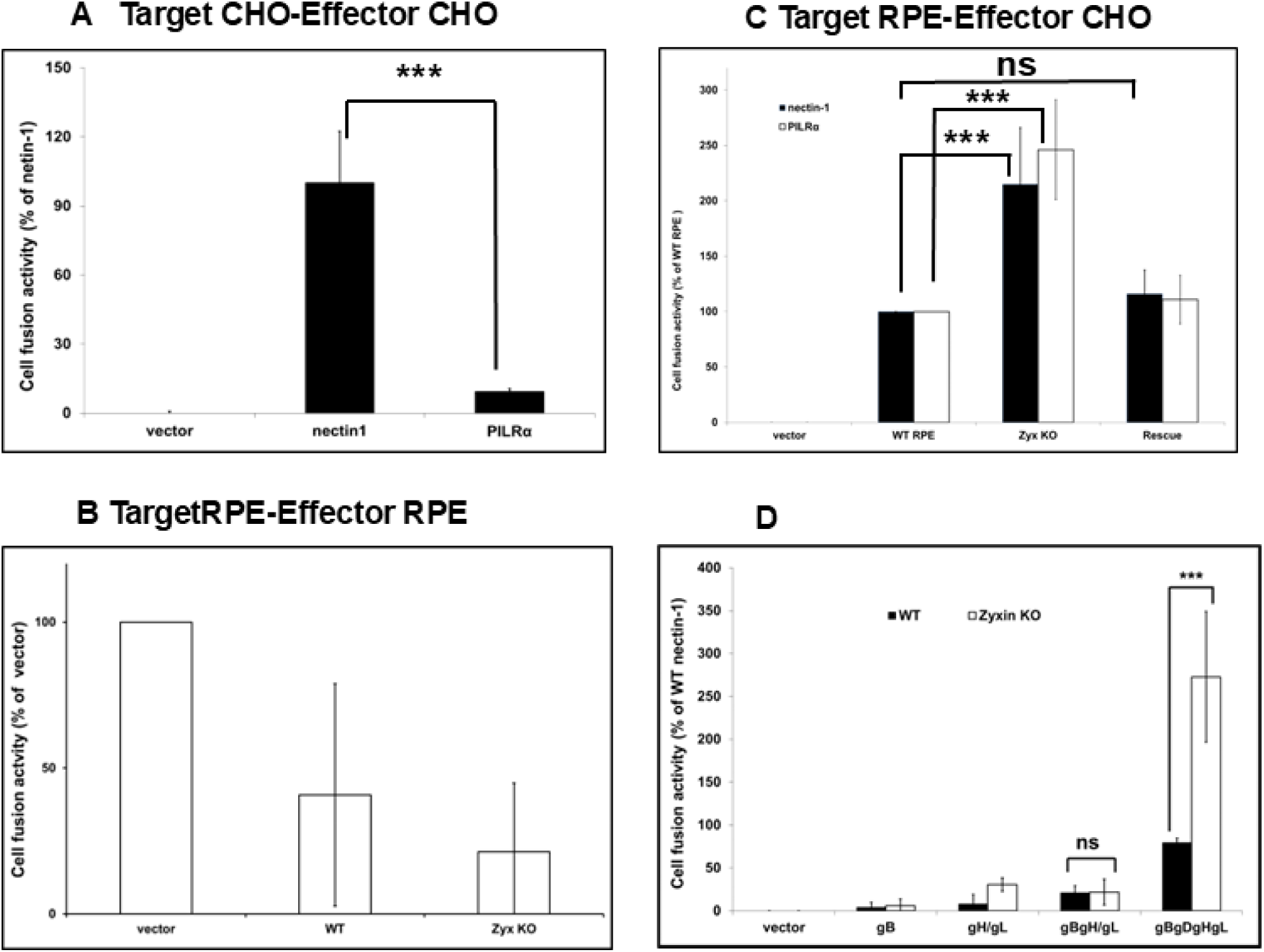
Cell-cell fusion by HSV-1 glycoproteins is enhanced in the absence of zyxin. Target CHO-K1 (A) or RPE (B and C) cells expressing luciferase under the control of the T7 promoter and either nectin-1 or PILRα (or an empty vector control) were incubated together with effector CHO-K1 (A and C) or RPE cells (B) expressing T7 polymerase and four HSV-1 entry proteins (gB, gD, gH, gL). (D). The cell fusion was done as like Fig.1 A-C, CHO-K1 effector cells expressing select groups of HSV-1 glycoproteins and target RPE cells expressing PILRα. Luciferase activity was measured as an indicator of cell-cell fusion. Backgrounds from empty vector controls were subtracted and data were normalized to the fusion activity observed with nectin-1. Results represent the means and standard deviations from three independent experiments. Error bars are standard deviations. Not significant (ns; P > 0.05) and asterisks (P < 0.01) are based on Paired Samples t Test.

Given the unprecedented high level of fusion observed with PILRα in zyxin-KO RPE cells (Fig. 1C), we investigated whether fusion between CHO-K1 effector and zyxin-KO RPE target cells could bypass the requirement for gD. Our results demonstrated that CHO-K1 effector cells required all four glycoproteins (gB, gD, gH, and gL) for fusion with PILRα-expressing RPE targets (Fig. 1D). These findings confirm that the hyperfusogenic fusion activity associated with PILRα remains gD-dependent, consistent with previous studies (26, 28).

### Cell fusion activity of PRV, hPIV5, and VSV glycoproteins

PRV uses orthologs of the HSV-1 glycoproteins to enter cells. Therefore, to reaffirm that removal of zyxin from target cells increases fusion, the PRV glycoproteins (gB, gD, gH, and gL) were tested in the cell-cell fusion assay with nectin-1 target cells. The human parainfluenza virus 5 (hPIV5) glycoproteins (HN and F), and the vesicular stomatitis virus (VSV) glycoprotein (G), were included to determine if the increased fusion observed in the absence of zyxin was specific to herpesviruses. RPE cells natively express sialylated host cell receptors (hPIV5 receptor) and low-density lipoprotein receptor (LDLR; VSV receptor), so the hPIV5 and VSV-target cells were transfected with a T7-driven luciferase plasmid and an empty vector. Increased fusion with zyxin-KO RPE target cells was observed with all viral glycoproteins. Relative to WT RPE target cells, fusion activity with zyxin-KO cells reached 247% for HN and F (hPIV5), 306% for G (VSV), and 170% for the PRV glycoproteins (*P* < 0.01) (Fig. 2A). Fusion activity of the rescue cells was between that of the WT and KO cells. These findings indicate that zyxin possess broad antiviral properties.

**Fig. 2.**
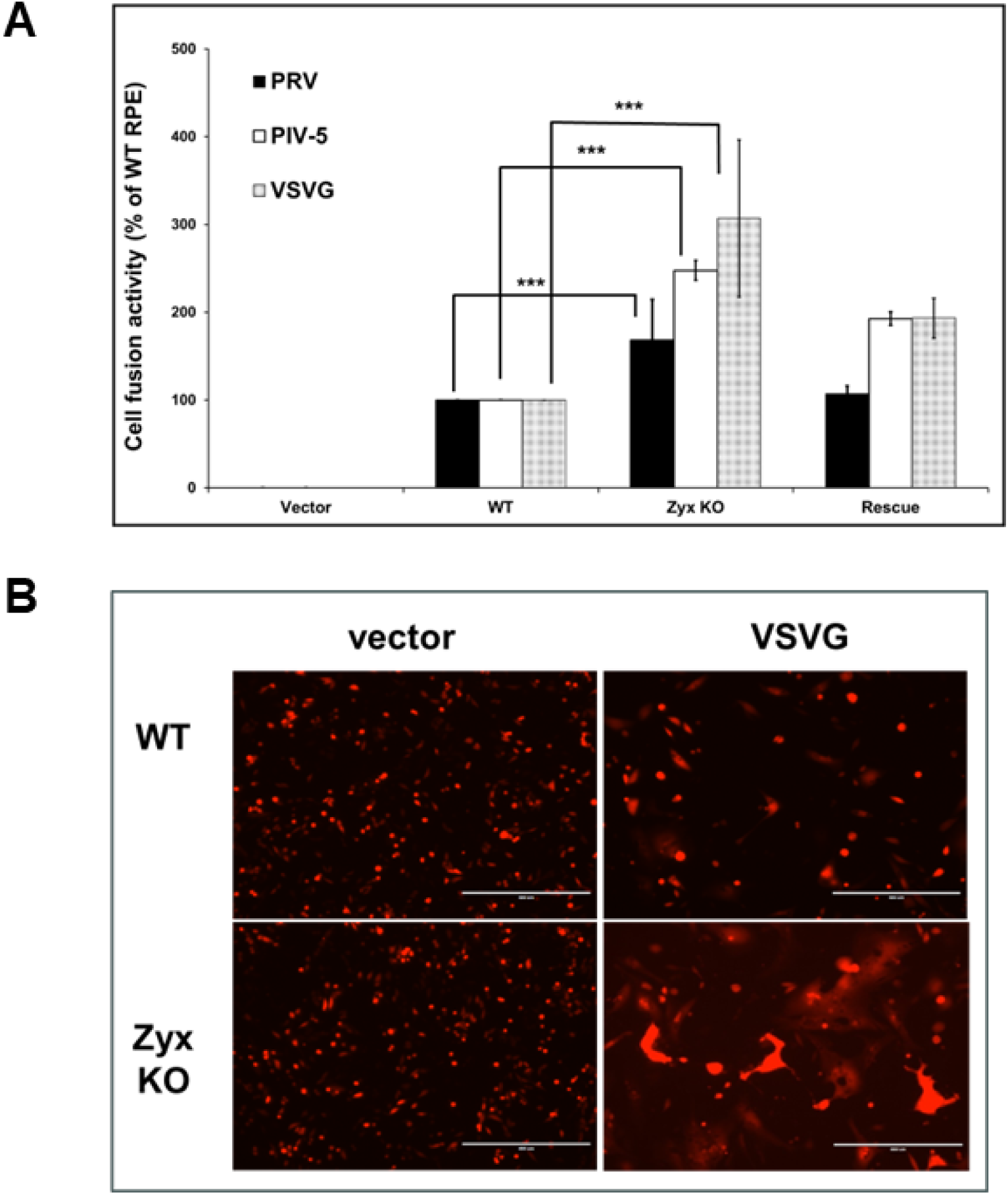
Cell-cell fusion by PRV, hPIV-5, and VSV glycoproteins is enhanced in the absence of zyxin. (A) Cell-cell fusion assays were performed as in figure 1 using effector CHO-K1 cells expressing either hPIV-5 F and HN, VSV G, or PRV gB, gD, gH, and gL. Target cells were WT, zyxin-KO, or rescue RPE cells expressing Nectin-1 (PRV) or transfected with an empty vector (hPIV-5 and VSV). Each bar shows the mean and standard deviation of at least three independent experiments. \*\*\**P* < 0.01 (Paired Samples Test). (B) Visualization of enhanced cell-cell fusion by VSV-G. CHO-K1 effector cells expressing VSV-G and RFP (or RFP alone) were co-cultured with target WT or zxyin KO RPE cells expressing GFP. Images were captured by fluoresce microscopy after 24 hrs.

The cell-cell fusion data obtained up this point provided a general assessment of fusion activity. To supplement these results, we next imaged the cells by fluorescence microscopy. CHO-K1 effector cells were co-transfected with an red-fluorescent protein (RFP) expression plasmid and either a VSV-G expression plasmid or an empty vector. The cells were then co-cultured with target WT or zyxin-KO RPE cells and imaged after 24 hours. Although fusion was evident throughout the WT and KO cell cultures, large syncytia were prominent in the KO cells consistent with increased fusion activity. In contrast, RPE effector cells co-transfected with the empty vector showed no detectable differences in fusion activity (Fig. 2B). These results added support to the findings that RPE cells lacking zyxin possessed increased susceptibility to viral glycoprotein mediated fusion.

### HSV-1 infection in WT and zyxin-KO RPE cells

We previously reported that HSV-1 and PRV produce enlarged plaques on RPE cells in the absence of zyxin, with the resulting viral titers nevertheless being equivalent between those produced on WT and zyxin-KO RPE cells (24).

These findings were verified here (Fig. 3A, C). This phenotype could result from either the virus spreading more rapidly between zyxin-KO cells, or by zyxin exerting its antiviral effect by compromising virion infectivity. To test the latter, HSV-1 stocks were produced on WT and zyxin-KO RPE cells and the harvested stocks were used to infect Vero cells. Plaque sizes were similar between the two stocks indicating that the infectivity of HSV-1 virions was not impaired by zyxin (Fig. 3B, D).

**Fig. 3.**
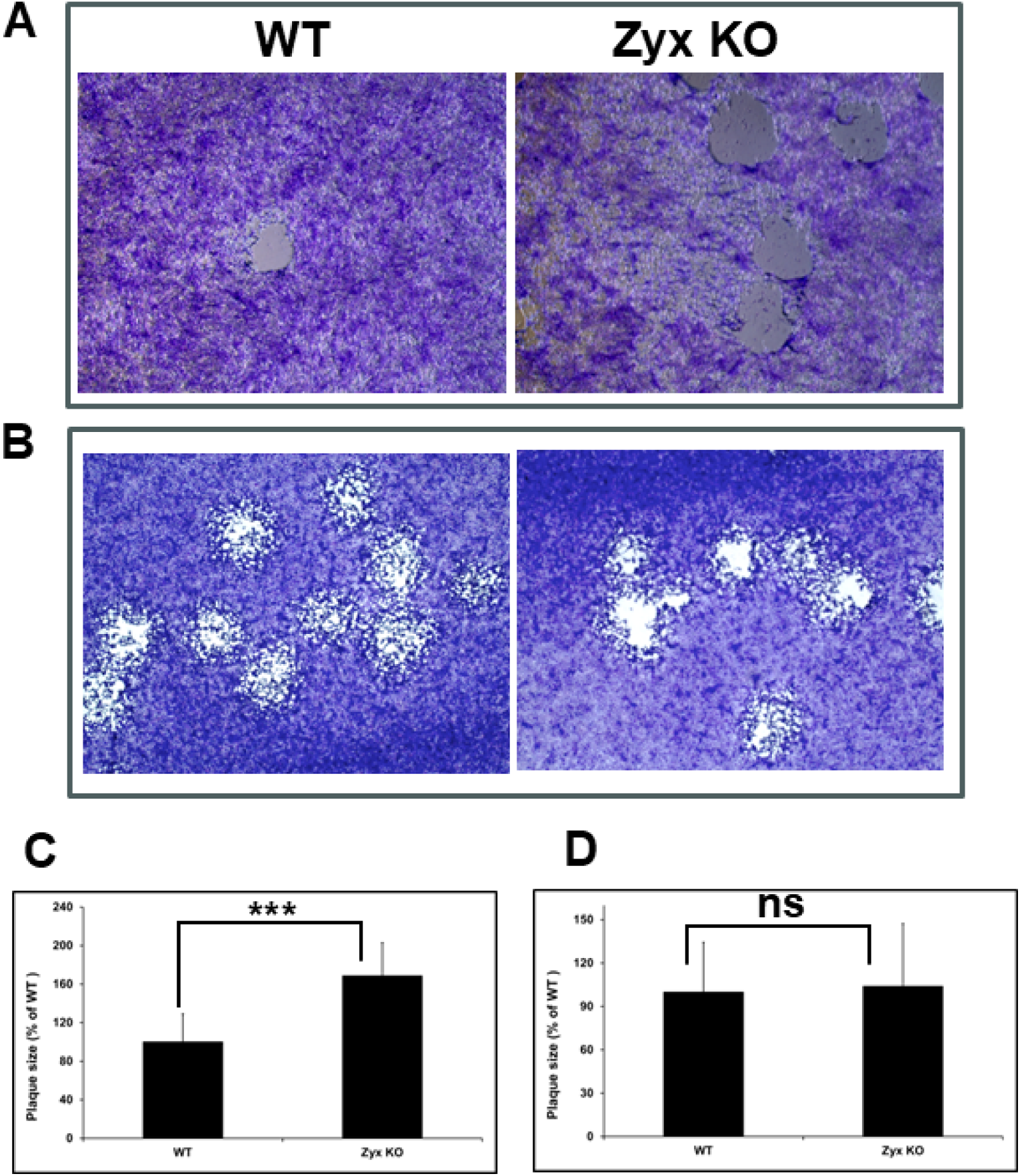
HSV-1 produces large plaques on zyxin-KO RPE cells. (A) WT or zyxin-KO RPE cells were infected with HSV1-GS3217 (MOI 0.01) and Giemsa stained at 3 d.p.i. (B) Viruses harvested from infected WT and zyxin-KO cells were used to infect WT RPE, Zyxin KO or Vero cells (200 pfu/well) and Giemsa stained at 3 d.p.i. Plaque images were captured under 4× magnification. (C and D) The radius of HSV-1 plaques was measured for a minimum of 50 plaques per sample and plotted as a percentage of the WT RPE cell sample. Error bars represent standard deviations. Asterisks indicate a significant difference in plaque size between the WT and zyxin-KO RPE cells (Mann-Whitney U test, *P* ≤ 0.01. ns, *P* >0.05).

### Plaque morphology of VSV and hHIV-3 on WT and zyxin-KO RPE cells

Given that hPIV5 and VSV glycoproteins also exhibited hyperfusogenic activity with zyxin-KO target cells (Fig. 2), we investigated whether plaque formation by paramyxovirus and rhabdovirus was affected. For this study, we used VSV and hPIV3, as hPIV5 was not readily available. In both cases, plaque sizes were substantially larger on zyxin-KO cells than on WT RPE cells (3.4x for VSV and 4.3x for hPIV3; *P* < 0.01) (Fig. 4).

**Fig. 4.**
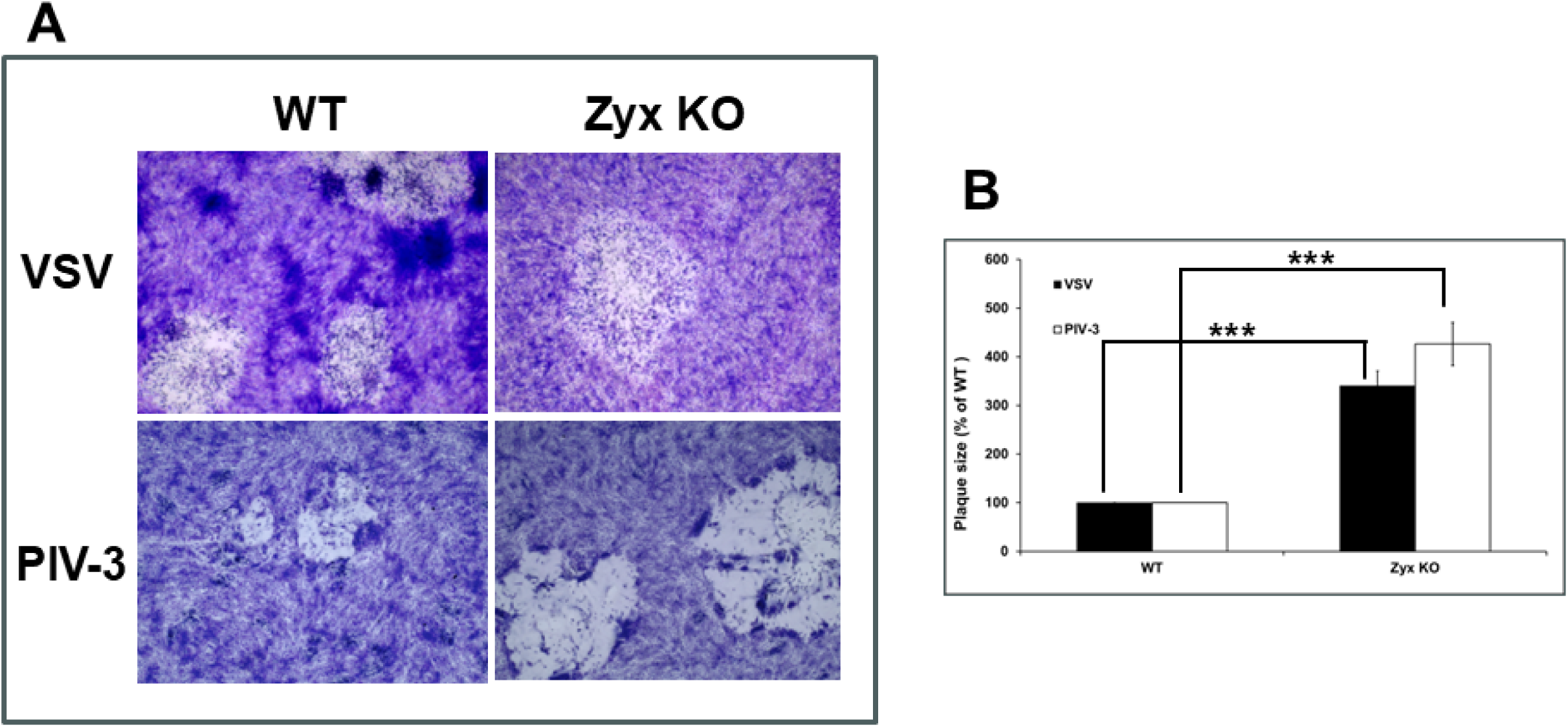
VSV and hPIV-3 produce large plaques on zyxin-KO cells. WT and zyxin-KO RPE cells were infected with the indicated viruses. (A) Cells were stained with Giemsa at 3 d.p.i. (B) Plaque sizes were analyzed and plotted as in figure 3. Asterisks indicate a significant difference in plaque size between the WT RPE and zyxin-KO cells (Mann-Whitney U test, *P* ≤ 0.01).

### Entry rate of HSV-1 and PRV into WT RPE cells and zyxin-KO cells

To determine if the observed increases in fusion (Fig. 1) and plaque sizes (Fig. 3) in zyxin-KO cells was associated with an increased rate of HSV-1 entry, the penetration kinetics of HSV-1 was measured in WT and KO RPE cells (29–31) (Fig. 5). HSV-1 was added to cell monolayers at 4°C for 1 hour to allow binding, then at 37°C for 10–60 minutes to allow entry, followed by acid inactivation of remaining extracellular virions. PRV was included for comparison, and duplicate samples rinsed with PBS instead of citric acid served as controls. After three days, plaques became evident and were counted to quantify productive viral entry events. The rate of HSV-1 and PRV entry was increased in zyxin-KO RPE cells, but by 60 minutes entry rates for both viruses equalized between WT and zyxin-KO RPE cells.

**Fig. 5.**
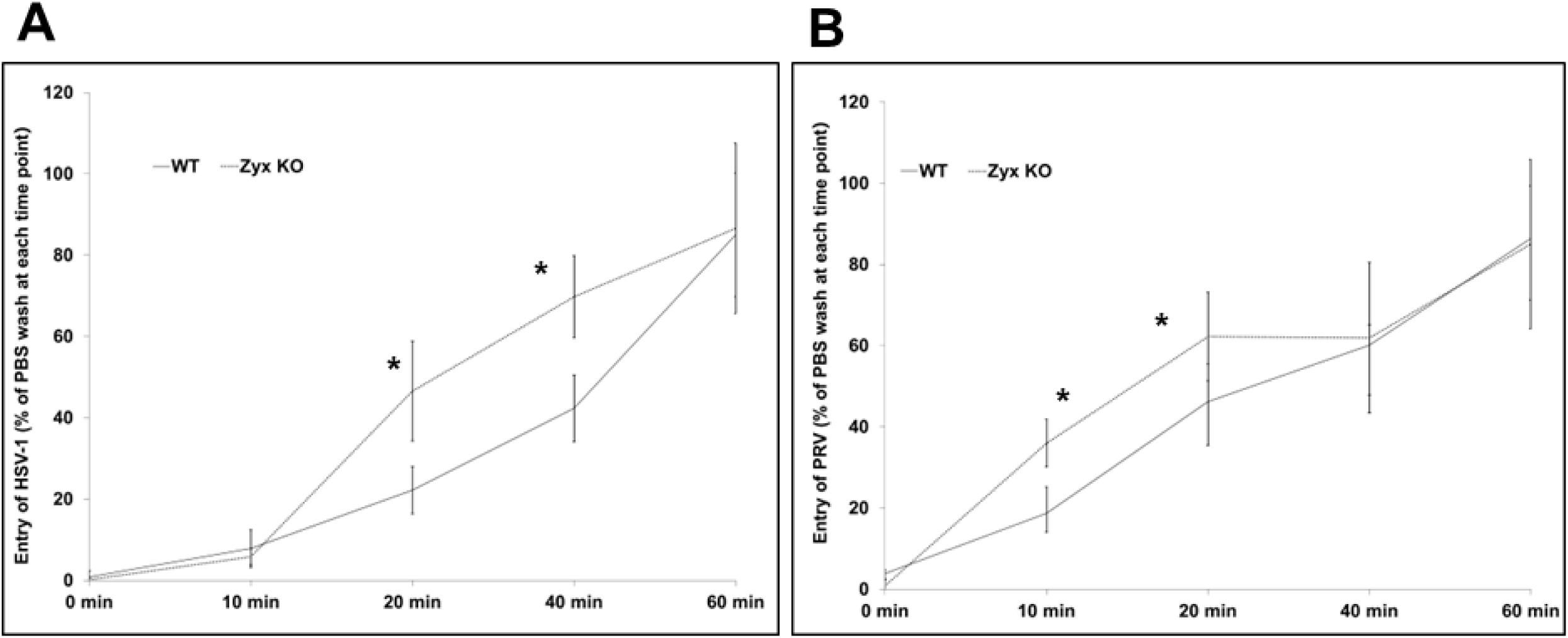
The rate of HSV-1 and PRV entry is increased in zyxin-KO cells. WT and KO RPE cells were incubated with the indicated virus at 4°C for 1 h before shifting to 37°C for 10 to 60 min. At the times indicated, the remaining extracellular virions were inactivated by citrate wash (or PBS as controls) and overlayed with methylcellulose for subsequent quantitation by plaque assay. Experiments were performed at least 2 times. The plaque counts are plotted as a percentage relative to the corresponding PBS wash control. Standard deviation is shown for each sample. \**P* < 0.01 (Paired Samples t Test).

### Bulk RNA sequencing of WT cells and zyxin-KO RPE cells

The cellular mechanisms by which zyxin constrains fusion and entry was explored by bulk RNA sequencing of uninfected WT and zyxin-KO RPE cells. Expression profiles for 58,396 genes were generated. Of these, 21,594 genes exhibited at least a 20% change in expression between the two samples, and 870 genes showed significant expression changes based on their log2-fold change (LFC) threshold. Based on these data, pathway analysis (https://metascape.org/) identified 18 genes that enrich into ontology several groups of interest, including regulation of membrane potential, herpes simplex virus 1 infection (CCL2 and ZNF14), NABA core matrisome (CCL2 and ZNF14), extracellular matrix organization (CTSK, SERPINE1 and TGFB2), cell-cell adhesion (RAC2 and TGFB2), anchoring fibril formation, and regulation of the MAPK cascade (ACKR3 and TGFB2). Eight of the latter are membrane proteins (ACKR3, SLC1A1, LDB2, A4GALT, GRIK5, LYPD6, OR2T8, TMEM171, and SLC26A11), of which ACKR3 serves as an HIV receptor (32, 33) and A4GALT is associated with HIV pathogenesis (32, 33). The remainder are non-membrane proteins (TGFB2, CTSK, SERPINE1, ANKRD1, FBXO32, MCM5, CCL2, CTSZ and NPTX1). The selected genes are indicated on a Volcano plot (Fig. 6). According to the analysis, 13 of these genes may be associated with viral infections, and approximately 50% may be related to endocytosis. We are particularly interested in membrane proteins, as they can play key roles in viral entry and membrane fusion. These proteins may influence fusion directly or indirectly by modulating membrane dynamics, receptor trafficking, or lipid raft integrity, processes that are often linked to endocytosis. Additionally, some non-membrane proteins, such as ANKRD1 and TGFB2, have been reported to be associated with endocytic pathways (34, 35). Further studies will be necessary to elucidate the precise mechanisms through which these proteins regulate viral infection and cell fusion.

**Fig. 6.**
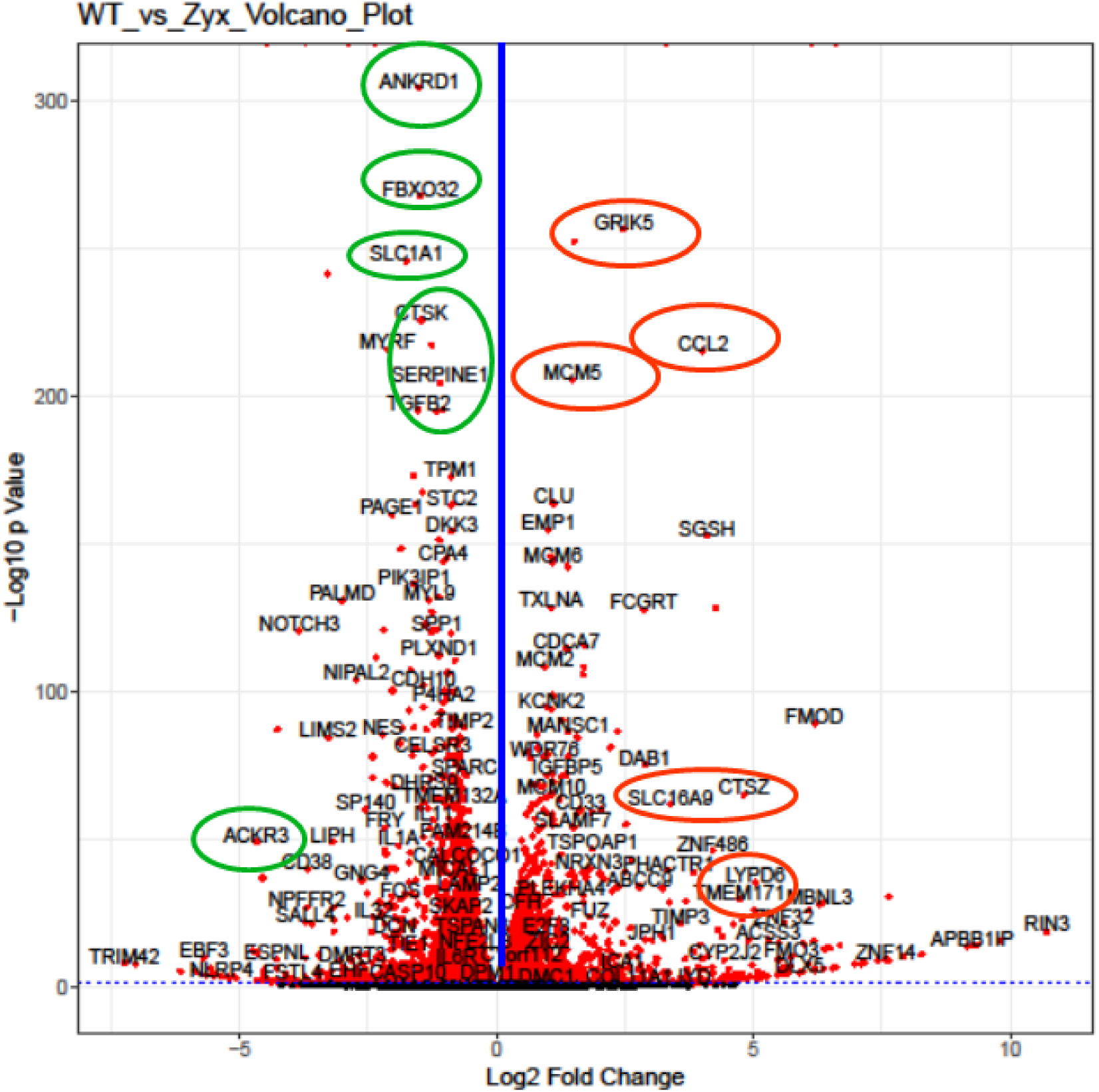
Volcano plot of differentially expressed genes identified between the WT RPE and zyxin knock out cells. The data points left of the blue line denote down-regulated expression, and the points on the right denote up-regulated expression. The circled genes are of interest (see text).

### Endocytosis is enhanced in zyxin-KO RPE cells

Enveloped viruses typically enter cells by fusing their envelope with the cell plasma membrane or an endosomal membrane, in the latter case after initial uptake. HSV-1 can enter cells by either pathway depending on cell type (36, 37). Because the RNAseq analysis indicated zyxin may modulate endocytosis, we monitored the internalization of fluorescently-coupled dextran (FITC-dextran) in WT and zyxin-KO RPE cells. FITC-dextran serves as a fluid-phase marker for uptake by endocytosis. RPE cells grown on coverslips were incubated with 1 mg/ml FITC-dextran for 1 hour at room temperature, followed by imaging with an EVOS Cell Imaging System. ImageJ analysis indicated a 53% increase in fluorescence intensity in zyxin-KO cells relative to WT RPE cells (*P* < 0.01) (Fig. 7A, B). To examine endocytosis further, cells were seeded in 96-well plates, incubated with FITC-dextran, and measured for fluorescence intensity using a PerkinElmer plate reader. The FITC signal in zyxin-KO cells was 190% of that in WT RPE cells (*P* < 0.01) (Fig. 7C). These findings implicate zyxin playing a role in restraining endocytosis and show that there is a correlation between increased endocytosis and increased viral fusion susceptibility.

**Fig. 7.**
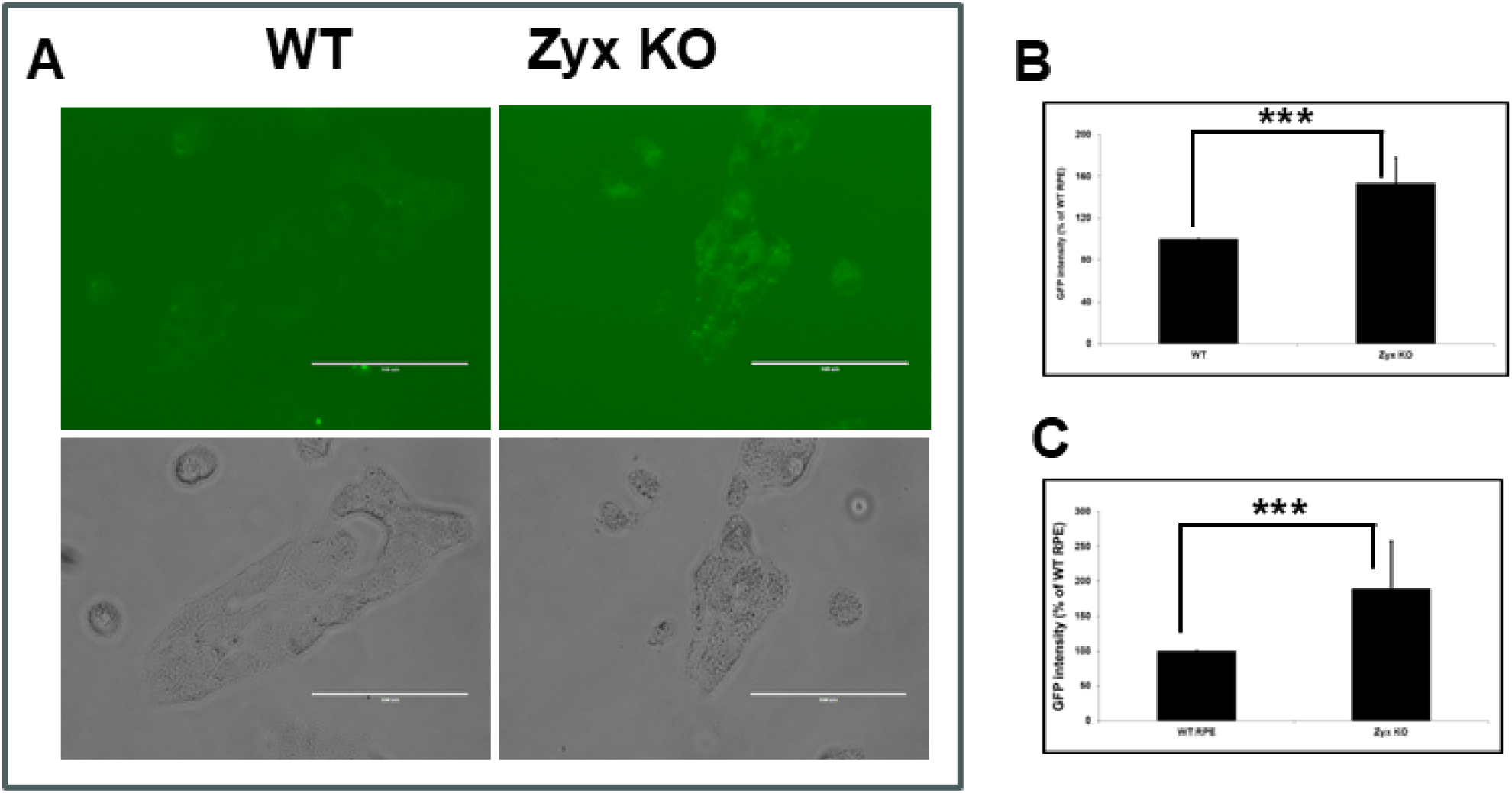
Endocytosis is upregulated in zyxin-KO cells. WT and zyxin-KO RPE cells were incubated with FITC-dextran to measure fluid-phase endocytic uptake. (A & B) Transmitted light and fluorescence emissions were imaged by microscopy under 40× magnification and plotted as a percentage of the WT RPE cells. (C) FITC-dextran uptake measured using a plate reader. All experiments were repeated a minimum of three times. Asterisks indicate a significant difference between the WT and zyxin-KO RPE cells (Mann-Whitney U test, *P* ≤ 0.01).

### Nectin-1 expression in WT and zyxin-KO RPE cells

The increased endocytosis observed in zyxin-KO RPE cells had us question if Nectin-1 expression on the cell surface was altered. Confluent WT and zyxin-KO RPE cells were stained with BV650-conjugated anti-Nectin-1 antibody (clone CK41), and the percentage of BV650-positive cells along with their median fluorescence intensities (MFI) were determined. The results showed that the fraction of cells expressing detectable cell surface levels of Nectin-1, and the MFI of the nectin-1-positive subpopulations, were increased in the absence of zyxin (153.2% in cell number and 148.3% in MFI; p < 0.001) (Fig. 8A). RNA sequencing revealed upregulation of nectin-1 and PILRα in zyxin-KO cells relative to WT RPE cells (*P* < 0.01 ) (Fig. 8B). LDLR (a receptor for VSV) in zyxin-KO cells was upregulated 33% in the zyxin-KO RPE cells. These results suggest that despite the increase in fluid-phase endocytosis associated with zyxin-KO cells, these cells have heightened expression of multiple cell receptors that are required for efficient fusion-based entry of several enveloped viruses.

**Fig. 8.**
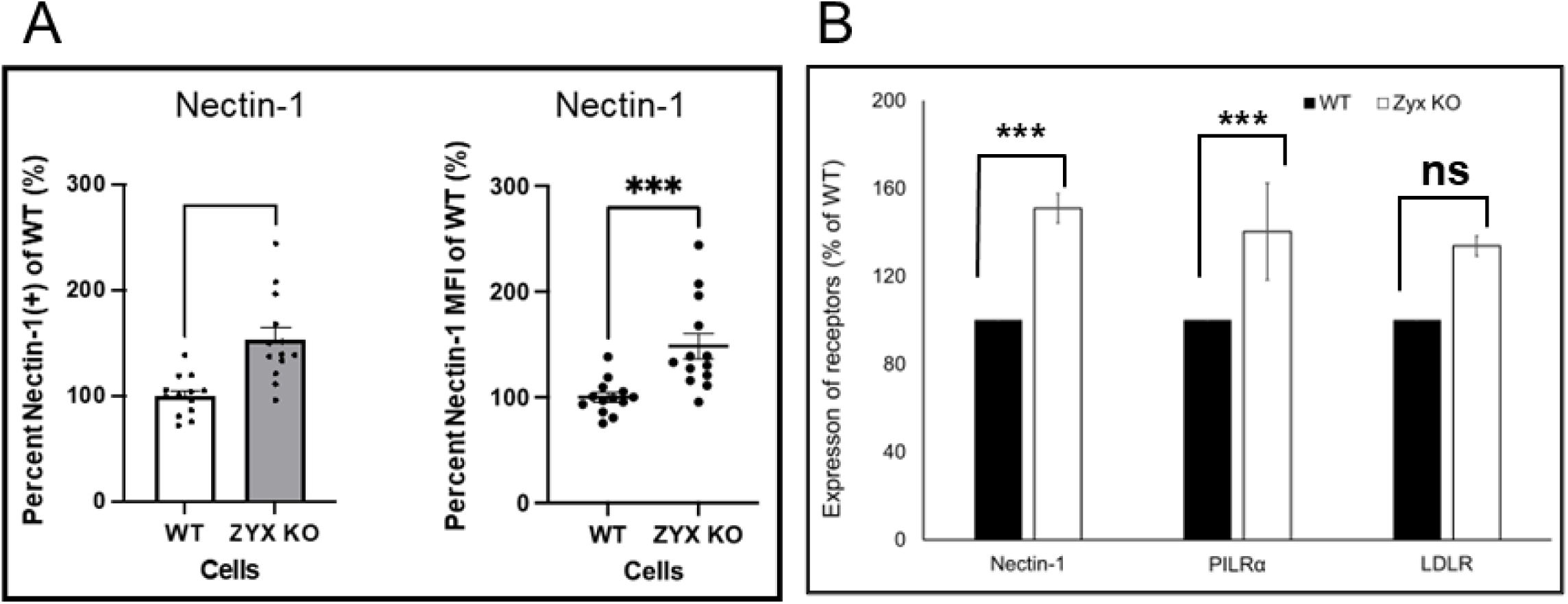
Expression of virus receptors is increased in zyxin-KO cells. (A) Cells were stained with mouse BV650-conjugated Nectin-1 antibody. Values are the percent BV650-positive measured by flow cytometry and are plotted as the mean of four independent experiments, each with internal triplicates. Error bars are SEM. Unpaired t-test *** p < 0.001. (B) RNA expression of nectin-1, PILRα, and LDLR based on differential expression analysis of RNA seq data. All data is normalized to WT RPE cells.

## DISCUSSION

Cellular factors constraining viral entry can antagonize infection at the initial step of infection. Zyxin is a multifunctional cytosolic protein that can directly modulate the cytoskeleton and engage in signal transduction to alter nuclear gene expression. Regarding the former, zyxin facilitates Cdc42 or WASP recruitment to cell surface receptors and plays key roles in actin polymerization and filopodium formation (19). Regarding the latter, zyxin interaction with the HPV-6 E6 protein mobilizes zyxin import into the nucleus (38). An example of the interplay of zyxin with cytoskeleton dynamics and nuclear signaling is seen in endothelial cells, where zyxin interacts with PAR-1 to coordinate thrombin signaling with actin remodeling. Zyxin depletion delays endothelial barrier restoration after thrombin exposure (20). In addition, zyxin acts as a facilitator of the innate immune response to viral infection by stabilizing the interactions between RIG-I (Retinoic acid-inducible gene I) and MAVS, promoting type I interferon production and enhancing the cell’s ability to combat viral pathogens (39).

We recently reported that zyxin impacts HSV-1 and PRV infection (24). In this study we initially aimed to investigate the role of zyxin in HSV-1 fusion. Using a qualitative luciferase cell-cell fusion assay (26, 27), we found that RPE cells do not support fusion on their own, even in the presence of nectin-1. However, fusion was observed when CHO-K1 was used as the effector cells and RPE was used as the target cells. Zyxin KO cells showed significantly higher fusion activity than WT RPE cells when the target cells expressed nectin-1 or PILRα (Fig. 1). However, fusion remained dependent on HSV glycoproteins gB, gD, gH, and gL (Fig. 1D). Perhaps most interestingly was the finding that enhanced fusion extended to PRV, hPIV5, and VSV glycoproteins (Fig. 2A), suggesting zyxin broadly inhibits viral fusion. Fluorescence microscopy confirmed these findings, revealing large syncytia in zyxin-KO cells (Fig. 2B). Consistent with increased fusion, zyxin-KO cells also supported larger viral plaques for HSV-1, PRV, VSV, and hPIV3 (Figs. 3–4), and HSV-1 and PRV initially entered zyxin KO cells faster than WT RPE cells (Fig. 5).

Transcriptome analysis focused our attention on endocytosis as a possible mechanism by which zyxin impacts viral fusion, which can influence viral entry and receptor availability. RNA-seq of WT and zyx-KO RPE cells revealed differences in expression of genes that encode membrane proteins or are associated with endocytic trafficking (Fig. 6). However, whereas all viruses initially engage cell-surface proteins at the plasma membrane, the viruses used in this study fuse at different sites. VZV uses clathrin-mediated endocytosis to initially enter cells and then fusion is triggered with the surrounding endosomal membrane, whereas parainfluenza viruses fuse at the cell surface (40). HSV can fuse either at the cell surface or after endocytosis depending on cell type (37, 41, 42) (36, 43). While these differences in virus entry do not immediately predict a shared role for zyxin during infection with these agents, increases in fluid-phase endocytosis were evident in the zyx-KO RPE cells. This finding suggests that zyxin antagonizes endocytosis and that enhanced endocytosis may promote the entry of distinct viruses that use different entry pathways to initiate infection (Fig. 7).

Despite increases in endocytic uptake that occurred in the absence of zyxin, cell surface expression of HSV-1 entry receptors like nectin-1 and PILRα was elevated (Fig. 8). Zyxin is reported to interface with the cell adhesion receptor, nectin-4, and such interactions could sequester cell surface proteins and reduce their accessibility to viruses. For example, the zyxin– nectin interaction facilitates zyxin localization to cell-cell adhesions (44). Nectin-1 acts as a major receptor for HSV-1 entry into RPE cells (45, 46).

Zyxin was initially identified in a screening for proteins interacting with viral tegument, a component not directly involved in cell–cell fusion. The effect of zyxin on cell-cell fusion across multiple virus families was unexpected. These findings suggest that zyxin, or zyxin-regulated proteins, suppress fusion and act as broad-spectrum antiviral factors. However, the mechanism behind this effect needs to be investigated further. Zyxin may act directly at the membrane-cytoskeleton interface to alter fusion pore formation or may act indirectly by altering endocytosis-related gene expression. While enhanced endocytosis may account for the larger plaque sizes observed in zyxin KO cells, the increased cell–cell fusion is likely independent of this pathway. Instead, the upregulated expression of some HSV-1 receptors in zyxin KO cells may partially explain enhanced fusion activity. Therefore, these effects of zyxin on viral fusion and entry may be attributed to increased receptor expression, enhanced endocytosis, and altered membrane potential regulation.

Although receptors for HSV-1 gD and gB are well characterized (47), other host factors that influence viral entry are unknown. Cells supporting plasma membrane fusion may express host proteins that functionally substitute for low-pH-triggered fusion in endocytic entry pathways. Zyxin-KO cells provide a powerful tool to identify unrecognized host factors that enhance fusion and entry. Future studies will focus on characterizing the genes identified in the RNA seq to elucidate their connection with zyxin as a viral entry restriction factor, potentially identifying new therapeutic targets for HSV-1 and other viruses.

## MATERIALS AND METHODS

### Cells and antibodies

African green monkey kidney Vero cells (American Type Culture Collection [ATCC], USA) and hTERT-RPE1 cells (RPE) were grown in Dulbecco modified Eagle medium (DMEM) supplemented with 10% fetal bovine serum (FBS) (Thermo Fisher Scientific, USA), penicillin, and streptomycin. The hTERT-RPE1 zyxin-KO cells and rescue cells (zyxin-KO + eGFP-zyxin) were additionally supplemented with 1 mM sodium pyruvate, and 4 µg/mL Blasticidin supplemented the rescue cell media. Chinese hamster ovary (CHO-K1) (ATCC, USA) cells were grown in Ham’s F12 medium supplemented with 10% FBS. All cell lines were maintained at 37°C in 5% CO_2_. BV650-conjugated Nectin-1 antibody clone CK41 (Invitrogen #BDB745384) was used at 1:100 diluted in MACS buffer (0.5% (wt/vol) BSA in D-PBS with 2mM EDTA).

### Plasmids

Plasmids encoding HSV-1 KOS strain gB (pQF439), gD (pQF440), gH (pQF441) and gL (pQF442) were previously described (48). PRV gB, gD and gL were cloned in pCAGGS and gH was cloned into pSG5 (Kindly provided by Dr. Barbara Klupp and Dr. Thomas Mettenleiter from Friedrich-Loeffler-Institut, Federal Research Institute for Animal Health, Germany). pT7EMCLuc plasmid encoding a firefly luciferase reporter gene under the control of the T7 promoter and pCAGT7 plasmid encoding T7 RNA polymerase (25, 49) were used in the fusion assay. The plasmids expressing human nectin-1 (pBG38) (kindly provided by Dr. Patricia Spear at Northwestern University) and human PILRα (pQF003) were described previously (26). hPIV5 pCAGGS HN and pCAGGS F were kindly provided by Dr. Sarah Connolly at DePaul University.

### Viruses

HSV1-GS3217 is a derivative of the HSV-1 F strain that encodes the CMV immediately early promoter driving the tdTomato fused to a nuclear localization signal (50). The CMV>NLS-tdTomato>pA cassette is inserted in the US5 (gJ) gene. VSV was from Dr. Robert Wagner (state institution to be consistent). Human PIV-3 (hPIV3) (influenza A/Japan/305/57) was from Dr. Rober Lamb at Northwestern University. PRV was a gift from Tamar Ben-Porat (Department of Microbiology Vanderbilt University School of Medicine, Nashville, Tennessee 37232 USA).

### Plaque size assay

All viruses used in this report form plaques on RPE cells that became visible two to three days post-infection. Contrast was increased by Giemsa staining prior to imaging by transmitted light microscopy using an EVOS Cell Imaging System. The average radius across a minimum of 50 plaques (two independent measurements per plaque) were collected for each sample. Using the average radius, the plaque area and the ratio of plaque size between WT and mutant viruses was determined, as described previously (29–31).

### Cell-cell fusion assays

To assess cell-cell fusion in cell populations, luciferase-based fusion assays were performed as previously described (25). Briefly, CHO-K1 cells and RPE cells (WT, zyxin-KO, and rescue) were seeded in 6-well plates overnight. For HSV-1 fusion, the effector CHO-K1 cells were co-transfected with five expression plasmids (400 ng each) encoding: T7 RNA polymerase, gB, gD, gH and gL (48), using 5 µl of Lipofectamine 2000 (Invitrogen, USA). For VSV and hPIV5 fusion, the effector CHO-K1 cells were transfected with pVSVG for VSV or pCAGGS hPIV5-HN and pCAGGS hPIV5-F for hPIV5. Target CHO-K1 or RPE cells were transfected with 400 ng of a plasmid encoding firefly luciferase under the control of the T7 promoter and 1.5 µg of a plasmid expressing either nectin-1, PILRα or an empty vector, using 5 µL of Lipofectamine 2000. After overnight transfection, the cells were detached with Versene and resuspended in 1.5 mL/well of F12 medium (CHO-K1 cells) or DMEM (RPE cells) supplemented with 10% FBS. Effector and target cells were mixed in a 1:1 ratio and re-plated in 96-well plates for 24 hours. Luciferase activity was quantified using a luciferase reporter assay system (Promega) and a Wallac-Victor luminometer (Perkin Elmer). To assess cell-cell fusion by microscopy**, t**arget and effector cells were transfected with fluorescent proteins. CHO-K1 cells were transfected with 400 ng of a plasmid encoding mcherry red fluorescent protein (RFP) along with 1500 ng of pVSVG or an empty vector. Transfected cells were co-cultured with target WT cells or zxyin KO RPE cells that had been transfected with 400 ng of plasmids encoding GFP. After coincubation for 24 h, pictures were captured using an EVOS Cell Imaging System at identical settings under ×200 magnification.

### FITC-dextran endocytosis assays

To assess endocytosis in individual cells, WT and zyxin-KO RPE cells were seeded in 6-well plates with a cover glass in each well and incubated overnight before adding 1 mg/ml FITC-dextran (Sigma, FD40S) dissolved in MEM (no glutamine and no phenol red, ThermoFisher) for 1 hour at room temperature. Fluorescence emissions were captured with an EVOS Cell Imaging System at identical settings under 40× magnification, quantified with Image J, and expressed as a percentage of the WT RPE cells. To assess endocytosis in cell populations, cells were seeded in 96 well black/clear plates and incubated overnight. FITC-dextran dissolved in MEM (1 mg/ml) was added for 1 hour at room temperature and the cells were then gently washed and the FITC signals were measured using PerkinElmer plate readers at an excitation wavelength of 480 nm and an emission wavelength of 530 nm. More green fluorescence signal indicates higher levels of endocytosis, as it reflects greater internalization of FITC-dextran by the cells.

### Viral titers and the virus release assay

WT or zyxin-KO RPE cells were infected with HSV1-GS3217, PRV, hPIV3 or VSV. The cells and supernatants were harvested separately, then centrifuged for 10 minutes at 13,000 rpm, and the cellular debris was merged with the cells from the same well and subjected to three freeze-thaw cycles. The harvested samples were tittered on Vero cells and stained with Giemsa 3 days postinfection.

### Determination of cell surface nectin-1 expression

Confluent WT or zyxin-KO RPE cells were lifted from flat-bottom 96-well plates with 0.2% (wt/vol) tetrasodium EDTA in PBS (Gibco Versene, #15040066). Following the detachment period, 10% serum media was added to wells and the cell mixture was transferred into round-bottom plates. Cells were washed once with MACS buffer prior to incubation with BV650-conjugated Nectin-1 antibody clone CK41 (Invitrogen #BDB745384) diluted 1:100 in MACS buffer at RT for 30 mins in the dark. Cells were washed with MACS buffer and resuspended in 1% paraformaldehyde in D-PBS. The Attune NxT Acoustic Focusing flow cytometer (ThermoFisher) and Attune Nxt Software were used to capture exactly 5000 events per sample. All data was analyzed on FlowJo v10.8.1. Events were gated to exclude debris and doublets. The BL-1 (GFP) channel was used to detect and exclude autofluorescent cells. Only cells that fell above the threshold for the Brilliant Violet 650 (VL-4) were considered positive for Nectin-1. The FlowJo software was used to determine the median fluorescence intensity (MFI) of the Nectin-1-positive population. In some cases values were normalized against WT intensities.

### Virus penetration kinetics

Measurements of viral entry into WT and zyxin-KO RPE cells were made by penetration assay as described previously (29, 51). HSV-1 and PRV were diluted in PBS and incubated with the WT or zyxin-KO RPE monolayers in 6 well plates for 1 h on ice. Cells were washed 3 times with cold PBS and then 1 mL/well of medium prewarmed to 37°C was added (time 0). Cells were incubated at 37°C for up to 60 minutes. At the indicated time points, cells were treated for one minute with either citrate buffer (135 mM NaCl, 10 mM KCl, 40 mM citric acid, pH 3.0) or PBS as a control. After the treatment, the cells were rinsed carefully five times with PBS. The monolayers were overlaid with 0.5% methylcellulose in DMEM with 1% heat-inactivated serum (Sigma-Aldrich, USA) and incubated at 37°C for 3 days. Plaques were visualized by Giemsa staining and counted under light microcopy.

### Bulk RNA sequencing

Cell pellets from sub-confluent 6-well plates were prepared in triplicates. RNA was extracted using the RNeasy Plus mini kit (Qiagen). Library preparation and sequencing were conducted by Northwestern NUSeq (Genomic Services), with each sample sequenced to a depth of 20 million reads.

### Statistical analysis

Statistical comparison of the plaque areas was performed with a Mann-Whitney U test using SPSS (version 25). The standard two-tailed Student’s *t*-tests were implemented for direct pairwise comparisons in entry and fusion experiments. Error bars indicate standard errors between experiments. *P* values represent the following: *, *P <* 0.05; **, *P <* 0.01; and ***, *P <* 0.001, ns, not significantly difference. Analyses were performed using IBM SPSS statistics version 25 for Windows (IBM Corp., Armonk, NY) and SAS 9.4 (SAS Institute, Cary, NC).

## ACKNOWLEDGMENTS

We thank Dr. Yasushi Kawaguchi for providing the parental HSV-1 strain F BAC clone. We thank Nanette Susmarski for excellent technical assistance and members of the Longnecker laboratory for their help in these studies. R.L. is the Dan and Bertha Spear Research Professor in Microbiology-Immunology. G.S. declares financial interests in Thyreos Inc and EG427. Research reported in this publication was supported by the National Institute of Allergy and Infectious Disease (NIAID) of the National Institutes of Health under grant number AI148478 and AI148780 The content is solely the responsibility of the authors and does not necessarily represent the official views of the National Institutes of Health. This manuscript is the result of funding in whole by the National Institutes of Health (NIH). It is subject to the NIH Public Access Policy. Through acceptance of this federal funding, NIH has been given a right to make this manuscript publicly available in PubMed Central upon the Official Date of Publication, as defined by NIH.

